# Effects of age and sex on dendritic D2 autoreceptor inhibition in substantia nigra dopamine neurons

**DOI:** 10.1101/2022.07.10.498507

**Authors:** Eva Troyano-Rodriguez, Kylie Handa, Sarah Y Branch, Michael J Beckstead

## Abstract

Substantia nigra pars compacta (SNc) dopamine neurons are required for voluntary movement and reward learning, and advanced age is associated with motor and cognitive decline. In the midbrain, D2-type autoreceptors located on dendrodendritic synapses between dopamine neurons control cell firing through G protein-activated potassium (GIRK) channels. We previously showed that aging disrupts dopamine neuron pacemaker firing in mice, but only in males. Here we show that D2-receptor inhibitory postsynaptic currents (D2-IPSCs) in aged male mice are moderately smaller compared to young males as well as females, regardless of age. Local application of dopamine revealed a reduction in the amplitude of the D2-receptor currents in old males compared to young, pointing to a postsynaptic mechanism that could not be explained by impairment of the GIRK channels or degeneration of the dendritic arbor. Kinetic analysis showed no differences in D2-IPSCs in old versus young mice or between sexes. Potentiation of D2-IPSCs by corticotropin releasing factor (CRF) is also conserved in aging, indicating preservation of plasticity mechanisms. These findings have implications for understanding dopamine transmission in aging in both sexes and could explain in part the increased susceptibility of males to SNc degeneration of dopamine neurons in neurodegenerative disorders such as Parkinson’s disease (PD).

## Introduction

Dopamine neurons of the substantia nigra pars compacta (SNc) are integral to motor function and reward learning (Cohen et al., 2009; Schultz, 2007; Yin et al., 2008). Dopaminergic behaviors decline with advancing age, with movement deficits and apathy or depression commonly observed in the elderly population (Seidler et al., 2010; Taylor et al., 2022). Moreover, aging is the single largest risk factor for development of Parkinson’s disease (PD, reviewed in Reeve et al. (2014), where degeneration of SNc dopamine cells is the main hallmark of the disease and is responsible for the motor symptomatology (Hornykiewicz, 1975). While modest dopamine cell death is thought to occur with normal age (Haycock et al., 2003; Kubis et al., 2000; McGeer et al., 1988; Wang et al., 2022), it is insufficient to explain the ~60% loss in extracellular dopamine that has been reported in the striatum (Colebrooke et al., 2006), suggesting a contribution from declining physiological and/or morphological factors. Indeed, we have previously reported that aging affects pacemaker firing frequency and regularity in dopamine neurons from male mice (Branch et al., 2014), while firing in female mice is largely preserved (Howell et al., 2020). We have also observed a striking decrease in nimodipine-sensitive (presumably L-type) calcium channel currents in dopamine neurons from aged males, while currents through hyperpolarization-activated cyclic nucleotide–gated (HCN) channels and small-conductance calcium-activated potassium (SK) channels are unaltered. It is less clear how age affects other channels and receptors, and no study to date has explored individual synaptic inputs to dopamine neurons across the lifespan.

Dopamine neurons in the midbrain communicate with each other via dendrodendritic synapses (Groves & Linder, 1983), which express D2 type autoreceptors that sense the local release of dopamine (Lebowitz et al., 2021). Evoking local dopamine release through electrical or optogenetic techniques produces a D2 receptor-mediated inhibitory postsynaptic current (D2-IPSC) that is action potential- and calcium-dependent (Beckstead et al., 2004; Tschumi et al., 2022). This form of dendritic neurotransmission proceeds through D2-autoreceptor activation of Gi/o proteins and activation of type 2 inhibitory G-protein-activated inwardly rectifying potassium (GIRK) channels (Beckstead et al., 2004; Davila et al., 2003; Neve et al., 2004). The resulting inhibitory conductance hyperpolarizes the cell (Kim et al., 1995) and strongly reduces firing rate (Pucak & Grace, 1994). D2-IPSCs are also susceptible to multiple forms of plasticity and are regulated by different neuropeptides and neuromodulators (Beckstead et al., 2007; Beckstead et al., 2009; Beckstead & Williams, 2007; Piccart et al., 2015; Tschumi et al., 2022). For example, the stress related-molecule corticotropin releasing factor (CRF) transiently increases the amplitude of both D2- and GABA_B_ receptor-mediated currents in a postsynaptic manner consistent with activity at the level of the GIRK channel (Beckstead et al., 2009; Nimitvilai et al., 2012; Piccart et al., 2015; Piccart et al., 2019).

The aim of the present work is to delineate the extent to which pre- and postsynaptic components of DA neurotransmission in SNc are affected by advancing age in both sexes using mouse brain slices. We hypothesized that DA signaling would be reduced both pre- and postsynaptically resulting in smaller and slower currents, and that this deficit in dopamine transmission could affect dopamine cell function in advanced age. We did observe a significant reduction in D2 autoreceptor currents in aged males that was postsynaptic and at the level of the D2 receptor. However, both the kinetics and CRF-induced plasticity of D2-IPSCs are preserved with advanced age, as is dendritic branching of individual dopamine neurons in both sexes. These results indicate that most aspects of dendritic dopamine neurotransmission are largely resilient to normal aging.

## Results

### D2-receptor inhibitory postsynaptic currents are decreased in aged male mice

We first acutely prepared horizontal midbrain slices containing the SNc from young (2-8 month) and old (20-25 month) C57Bl/6N mice of both sexes. We then performed whole cell voltage clamp electrophysiological recordings of SNc dopamine neurons (holding voltage −55 mV). In the presence of synaptic blockers, we then used trains of five electrical stimuli to elicit D2-IPSCs (**Fig. 1A**) over a predetermined range of stimulus intensities (0.02-0.05-0.1-0.15-0.2-0.25-0.30 mA). Generally, D2-IPSCs appeared to be similar in amplitude and kinetics across ages and sex (**Fig. 1B**). On average, cells from old males exhibited the lowest maximal amplitudes [**Fig. 1C;** young males: 68.07 ± 10.89 pA (n = 15); old males: 45.97 ± 5.64 (n = 16); young females: 54.50 ± 6.76 (n = 16); old females: 66.68 ± 8.21 (n = 19)]. A two-way ANOVA did not indicate main effects of sex or age (sex, F_1,62_ = 0.1945, p = 0.6607; age, F_1,62_ = 0.3745, p = 0.5428); however, there was a significant age-sex interaction (F_1,62_ = 4.480, p = 0.0383). We then normalized each cell to its maximum amplitude, as a difference in the shape of stimulus-response curves could indicate alterations in presynaptic sensitivity or dopamine release probability. Data indicated the expected increase in amplitude with stimulus intensity (**Fig. 1D**, F_6,356_ = 351.9; P < 0.0001), which plateaued at the highest stimulus intensities. However, no difference was observed between experimental groups (F_3,62_ = 0.2957; p = 0.8284), nor was there a group-stimulus interaction (F_18,356_ = 0.4626; p = 0.9718). This suggests that the modest differences in aged male mice were not accompanied by an observable decrease in the stimulus-response relationship across groups, arguing against presynaptic differences. An analysis of the kinetics of the D2-IPSCs indicated that the average normalized traces from each group were essentially superimposed (**Fig. 1E**). No significant differences were observed between groups (F_3,57_ = 0.1083, p = 0.9549). These results suggest that the amplitude and time course of D2-IPSCs in largely conserved across age and sex, and the subtle difference observed in peak amplitudes cannot be explained by altered IPSC kinetics or a shift in the stimulus-response relationship.

**Figure 1.**
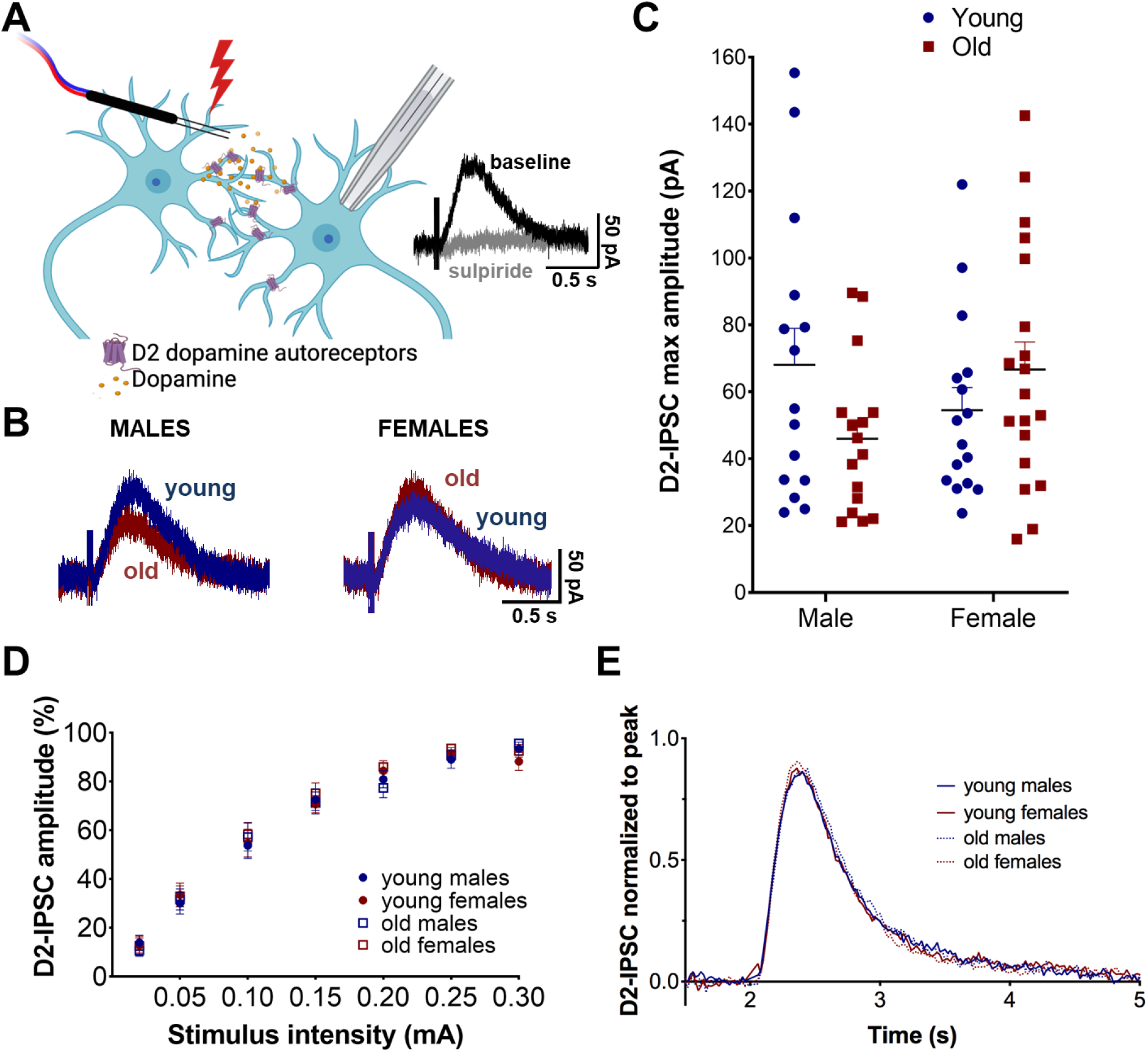
Effect of age and sex on D2 receptor inhibitory postsynaptic currents. **A)** Schematic diagram of stimulation of sulpiride-sensitive D2-IPSCs. **B)** Representative traces of D2-IPSCs from old versus young males and females. **C)** Maximum amplitude of D2-IPSCs for each individual cell for all groups. A significant sex x age interaction was found (*p* < 0.05). **D)** Normalized stimulus-response curves show no differences between groups. **E)** Kinetic analysis of D2-IPSCs amplitudes normalized to maximum outward current show no effect of age or sex.

### D2-receptor currents induced by exogenous dopamine are decreased in old male mice

Since we found that D2-receptor currents tended to be smaller in old males compared to young, we next investigated if this could be due to alterations in postsynaptic D2 signaling in aging males. We bath-applied a high concentration of dopamine (100 μM) for 7 minutes while holding the voltage at −55 mV. On average, peak amplitudes were larger in cells from young versus old males [**Fig. 2A, B** young: 217.77 ± 21.86 pA (n = 26), old: 178.39 ± 11.58 pA (n = 27)] but this effect was not statistically significant (unpaired two-tailed t-test, t_51_ = 1.609, p = 0.1138). Analysis of the normalized time course (**Fig. 2C)** beginning at the peak with 2-way ANOVA indicated a significant time x age interaction (F_13,540_ = 4.000, p < 0.0001) with main effects of time (F_13,540_ = 115.0, p < 0.0001) and age (F_1,45_ = 6.755, p = 0.0126). Posthoc analysis revealed a significant simple effect of age from 12-15 minutes following the drug application, indicating a possible decrease in desensitization following extended receptor activation.

**Figure 2.**
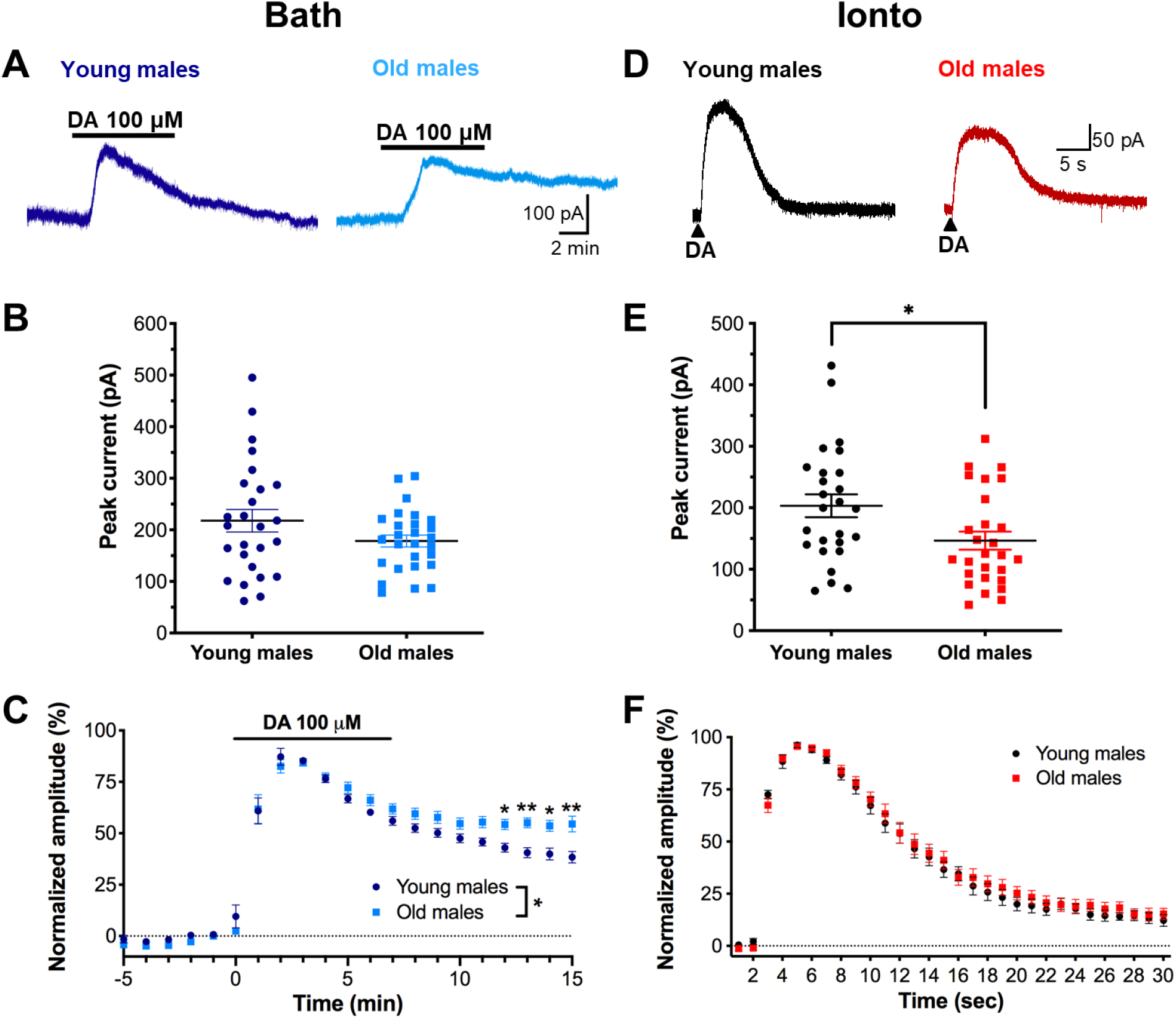
Outward currents in response to exogenous dopamine are reduced in old males. **A)** Representative traces of D2 receptor mediated currents elicited by bath-application of 100 μM dopamine (DA) for 7 min (horizontal line) from a young (left dark blue trace) and an old mouse (right light blue trace). **B)** Peak currents from individual cells. Old males tended to have smaller peak currents compared to young males, but this did not reach significance. **C)** Normalized time course of current elicited by bath-applied dopamine shows that responses in cells from old males do not desensitize to the same level after prolonged activation as young males. **D)** Representative traces from dopamine currents from young (left trace, black) and old males (right trace, red) when dopamine was applied by iontophoresis (Ionto). **E)** Peak currents from individual cells shows significantly smaller currents in cells from old males. **F)** Normalized time course of currents elicited by iontophoresis showed no difference between old and young males.

Since bath application of dopamine occurs slowly and peak amplitude occurs at a time that desensitization is starting to take hold, we repeated this experiment with rapid, local activation of dopamine receptors. We used iontophoresis to rapidly apply dopamine (1M in the pipette, ejection time: 1 second) near the soma as previously described (Beckstead & Williams, 2007; Sharpe et al., 2014). Analysis of the resulting D2-receptor current showed cells from old males have smaller currents compared to young male mice (**Fig. 2D, E**; young: 203.2 ± 18.58 pA (n = 26) and old: 146.4 ± 14.71 pA (n = 27); unpaired t-test, t_51_ = 2.404, p = 0.0199). When the currents were normalized to their peak amplitude (**Fig. 2F**), there was no significant effect age x time interaction (F_25,1225_ = 0.3975, p = 0.9967) that would indicate a difference in kinetics. Taken together, these results indicate that rapid, local application of dopamine produces smaller amplitude currents in cells from aged males, and desensitization may also be reduced in aged males following prolonged receptor activation.

### GABA_B_ receptor-GIRK channel signaling is not reduced in aged male mice

GIRK channels are coupled to D2-receptors in dopamine neurons and are responsible for the bulk of the D2-IPSC (Beckstead et al., 2004). We next sought to determine the effect of age on GIRK channel activation mediated by a different receptor. GABA_B_ receptors are also linked to GIRK channels in dopamine neurons, and activation of these receptors induces a potassium conductance that is partially overlapping with D2 receptor-mediated activation (Beckstead et al., 2009; Beckstead et al., 2004; Lacey et al., 1988). We therefore recorded from dopamine neurons of SNc in slices of young versus old male mice and sequentially bath-applied two concentrations of the GABA_B_ agonist baclofen (3 and 30 μM) while in voltage clamp (**Fig. 3A**). No effects of age were observed when measuring the magnitude of the outward potassium current induced by either concentration of baclofen (3 μM, young: 229.33 ± 14.724 pA, n = 9; old: 216.55 ± 19.193 pA, n = 10; 30 μM, young: 318.125 ± 20.628 pA, n = 8; old: 293.3 ± 23.287 pA, n = 10; 2-way ANOVA, age: F_1,17_ = 0.4918, p = 0.4926; concentration: F_1,16_ = 110.7, p < 0.0001, concentration x age interaction, F_1,16_ = 0.5541, p = 0.4674; **Fig. 3B**). These results suggest that the general functionality of the GIRK channels is conserved in young versus old male mice and that the differences in D2-receptor inhibitory synaptic currents observed in aging males are likely explained by differences at the level of the D2 receptor.

**Figure 3.**
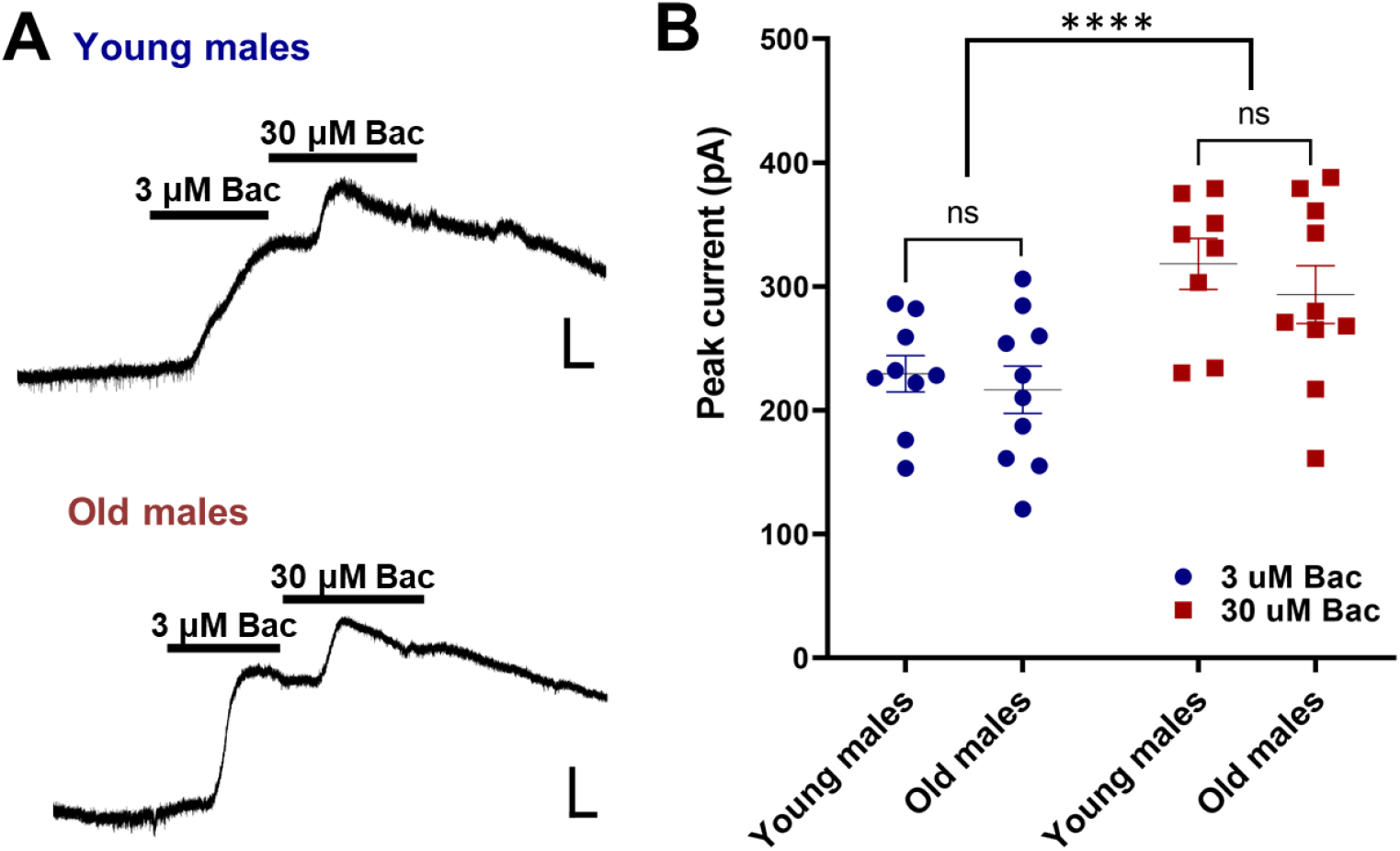
GABA_B_ currents are similar in dopamine neurons from old and young male mice. **A)** Representative traces of GABA_B_ receptor-mediated currents in young (upper trace) and old (lower trace) males. Two concentrations of the GABA_B_ agonist baclofen (3 and 30 μM) were applied sequentially to activate GIRK channels. **B)** Maximum amplitudes for individual cells at each concentration show no effect of age.

### D2-IPSC enhancement by the stress neuropeptide CRF is not affected by age

Corticotropin releasing factor (CRF) is a neuropeptide that has been widely implicated in both peripheral and central stress response. Along with its role in initializing the hypothalamic-pituitary-adrenal axis loop, CRF is also released in extrahypothalamic sites within the brain (Kelly & Fudge, 2018). CRF-positive axon terminals establish asymmetric (excitatory) synapses on VTA neurons that stain for tyrosine hydroxylase, a dopamine neuron marker (Tagliaferro & Morales, 2008). In dopamine neurons, CRF acts through CRF type 1 and type 2 receptors (CRF-1 and −2) (Sauvage & Steckler, 2001; Ungless et al., 2003) and has been reported to alter their firing rate and pattern (Korotkova et al., 2006; Lodge & Grace, 2005; Wanat et al., 2008). We have previously reported that bath application of CRF potentiates D2-IPSCs amplitude and slows time-to-peak (Beckstead et al., 2009), an effect blunted by both repeated psychostimulant administration and restraint stress. To determine if CRF-induced modulation of dendritic dopamine signaling is affected by aging, we recorded dopaminergic neurons from SNc and applied CRF at a concentration that potentiates D2-IPSCs (100 nM for 7-10 min; **Fig. 4A**).

**Figure 4.**
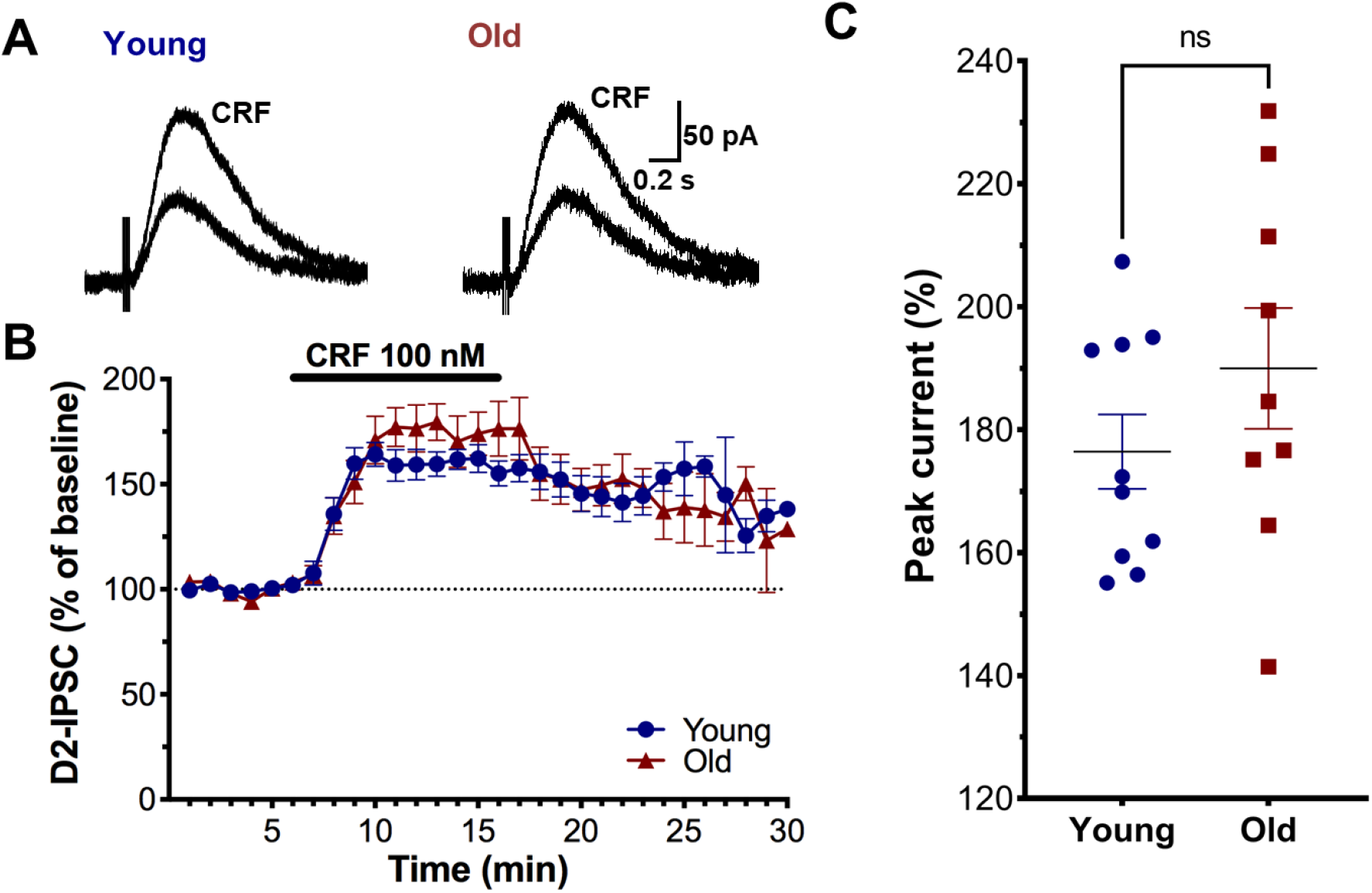
Corticotropin releasing factor (CRF)-induced plasticity is not affected by age. **A)** Traces from young and old mice depicting representative D2-IPSCs at baseline and after bath application of CRF (100 nM). **B)** Normalized time course of response to bath-application (horizontal bar) showed no effect of age. **C)** D2-receptor peak current achieved by CRF application in the recording bath for both studied groups. No differences were observed between young versus old mice.

Analysis of the time course (**Fig. 4B**) and peak enhancement (**Fig. 4C**) indicated that aging does not alter CRF-induced plasticity of D2 receptor synaptic currents (2-way ANOVA, age: F_1,17_ = 0.1685, p = 0.6865; time: F_29,382_ = 40.33, p <0.0001; time x age interaction F_29,382_ = 1.420, p = 0.0765; mean ± SEM, young: 176.4 ± 6.051 %, n = 10; old: 190.0 ± 9.828 %, n = 9; t_17_ = 1.201, p = 0.2462). This suggests that this form of D2-IPSC plasticity in dopaminergic neurons is preserved in aging.

### Aging does not alter somatodendritic morphology of SNc dopamine neurons

We next sought to determine if differences in somatodendritic morphology could be contributing to the alterations in D2 receptor-mediated currents that we observe in old males. Alterations in dendritic morphology have been previously described in dopamine neurons of SNc from adult (5.5-17 months of age) versus juvenile (2-4 weeks) female mice (Gantz et al., 2011) and by our group in both sexes of MitoPark mice, a genetic animal model PD (Lynch et al., 2018). Here, we measured dendritic morphology of SNc dopamine neurons located along the rostrocaudal and mediolateral axis of the SNc (DV −4.44 to −4.72 mm in the horizontal plane) by filling cells with biocytin included in the recording pipette, conjugating with streptavidin, and performing confocal imaging. Using the Sholl analysis method (Binley et al., 2014) we analyzed dendritic arbors between 20-400 μm from the soma (**Fig. 5A**). No differences were observed in dendritic branching between young and old mice (**Fig. 5B;** 2-way ANOVA, age: F_1,50_ = 0.07401, p = 0.7867; distance from soma: F_19, 950_ = 45.19, p <0.0001; age x distance from soma interaction F_19,950_ = 1.586, p = 0.0527). We further analyzed neurite morphology by quantifying multiple parameters. We first measured the total length of dendrites (young: 2,468 ± 248.8 μm, n = 31; old: 2,587 ± 284.4 μm, n = 22) and number (young: 21.74 ± 1.818, n = 31; old: 21.91 ± 2.114, n = 22; **Fig. 5E1-E2**) and no differences were observed between young and old mice (total length: t-test t_51_ = 0.3127, p = 0.7558; total number: t-test t_51_ = 0.05974, p = 0.9526). Since retraction of dendrites is more likely to happen towards the end of the branches, any subtle shrinkage would be reflected in a change of neurite length (or number) at some level of the dendritic arbor (branch order; **Fig. 5C**). We then took the mean length per branch for the different levels of the dendritic arbor as a key measure of shrinkage. No differences were observed between young and old groups (age: F_1,51_ = 0.1656, p = 0.6858; branch order F_4, 160_ = 3.874, p = 0.0050; branch order x age: F_4, 160_ = 0.4476, p = 0.7740; **Fig. 5D**). Another way to measure dendritic loss is by measuring the length and number of primary, inner and terminal branches (**Fig. 5C**). If somatodendritic morphology is affected by age, we would expect a decrease in the length and number of terminal branches with some inner branches possible becoming terminal (primary branches would be less likely to change). Again, no differences were observed in the length or number of primary branches (**Fig. 5F1-F2**, length primary; young: 430.3 ± 67.04 μm, n = 31; old: 383.1 ± 75.06 μm, n = 22; t_51_ = 0.4647, p = 0.6441; number primary; young: 4.323 ± 0.1934, n = 31; old: 4.136 ± 0.2573, n = 22; t_51_ = 0.5900, p = 0.5578), inner branches (**Fig. 5G1-G2,** length inner; mean ± SEM, young: 261.0 ± 50.73 μm, n = 31; old: 243.7 ± 70.01 μm, n = 22; t_51_ = 0.2051, p = 0.8383; number inner; young: 3.839 ± 0.4522, n = 31; old: 3.227 ± 0.5733, n = 22; t_51_ = 0.8469, p = 0.4010) or terminal branches (**Fig. 5H1-H2,** length terminal; mean ± SEM, young: 1684 ± 158.0 μm, n = 31; old: 1765 ± 165.5 μm, n = 22; t_51_ = 0.3454, p = 0.7312; number terminal; young: 13.19 ± 0.9418, n = 31; old: 13.45 ± 1.130, n = 22; t_51_ = 1.778, p = 0.8596). We conclude that SNc dopamine neuron morphology is preserved throughout the adult lifespan in both sexes, and that the reduced D2 receptor currents that we observed in old males are not due to shrinkage of the dendritic arbor.

**Figure 5.**
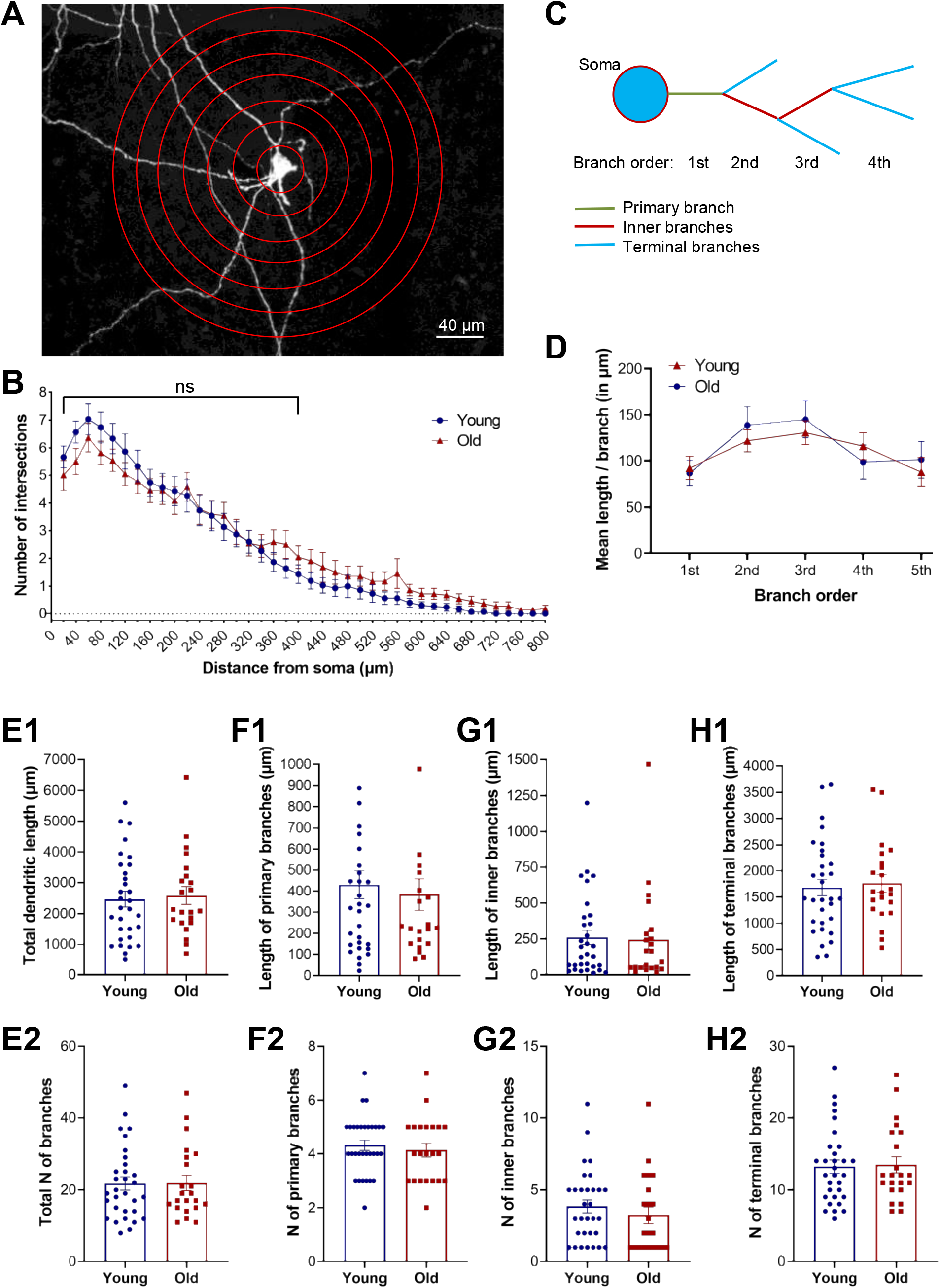
Somatodendritic morphology of SNc dopamine neurons is unaffected by age. **A)** representative confocal image of a labelled dopamine neuron with concentric circles at 20 μm increments used for the Sholl analysis. **B)** Average Sholl plot from old and young mice from the soma to 800 μm distance. Analysis was performed from 20-400 μm. No difference was observed between young and old mice. **C)** Schematic drawing of the soma and dendritic processes used for dendritic measurements. **D)** Mean length per branch for each branch order (from primary branches coming out of the soma to the 5^th^ level, after which the number of branches observed decreased considerably. **E1-2)** Total length and number of dendrites. Length and number of **F1-2)** primary, **G1-2)** inner and **H1-2)** terminal branches. No differences were observed between young and old mice.

## Discussion

This is the first report of the effects of aging on dopamine D2-autoreceptor transmission in SNc dopamine neurons from mice of both sexes. D2 autoreceptor-mediated synaptic currents were moderately reduced in dopamine neurons from aged males. While the stimulus-response curves were similar across all groups, iontophoretic application of dopamine near the vicinity of the recorded neuron produced smaller outward currents in aged males, suggesting a postsynaptic locus of action. This reduction in the maximal current amplitude did not appear to be caused by impairments at the level of the GIRK channels, as activation of GABA_B_ receptor currents with baclofen did not show an effect of age. Most other parameters were also unchanged by age, including CRF-induced enhancement of D2-IPSCs, the kinetics of the IPSC, and the quantification of dendritic arbor morphology of single dopamine neurons. As noted in our initial work describing aging in dopamine neurons (Branch et al., 2014), perhaps the most striking finding is the number of parameters that were *not* changed by age. However, the fact that many experiments yielded negative results gives us confidence that the observed decrements in D2 currents with age are robust and not due to experimental considerations, for instance as a phenomenological consequence of the brain slicing procedure.

In previous studies, we have found that dopamine neuron firing fidelity is compromised in aging only in males (Howell et al., 2020), and also observed a reduction in the nimodipine-sensitive (presumably L-type) calcium channel conductance with age (Branch et al., 2014). Our current results demonstrate that aging can affect synaptic input to dopamine neurons in addition to intrinsic conductances as evidenced by a reduction in D2 receptor-mediated inhibitory currents. This effect could potentially impact the precision and/or frequency of dopamine neuronal firing, along with other altered ionic conductances or synaptic inputs. We and others have previously shown a decrease in firing due to synaptically-evoked dopamine (Beckstead et al., 2004; Courtney et al., 2012). We have also long noted a slight but reliable increase in firing rate upon application of the D2 receptor antagonist sulpiride, suggesting modest tonic dopamine inhibition even in brain slices (data not shown). This is also consistent with recently published findings from other groups that have shown an inward current in response to sulpiride (Robinson et al., 2019) and an apparent rightward shift in firing rate histograms in the presence of sulpiride, although the latter was not directly compared (Shin et al., 2022). SNc dopamine neurons are notoriously sensitive to insults due to their high energy demands, large calcium oscillations, and handling of dopaminergic metabolites, and could be particularly susceptible to oxidative stress (Graves et al., 2020; Guzman et al., 2010; Surmeier et al., 2011). Age and sex are the two leading risk factors for development of Parkinson’s disease, with men exhibiting a 1.5-2.0 times higher incidence than women (Wooten, 2004). Increased firing in aged males due to reduced D2 receptor signaling could conceivably interact with other factors to initiate or accelerate processes associated with neurodegeneration due to normal aging and/or diseases such as Parkinson’s.

There are several potential explanations that could explain smaller D2-IPSC amplitudes in old males, which could include: 1) decreased presynaptic dopamine release, 2) altered postsynaptic D2 receptor activity, 3) altered GIRK channel activity, or 4) morphological changes, such as reduced dendritic branching or a decrease in synapse number. Although each of these factors does not preclude influences from the others, our findings are most consistent for effects at the level of the postsynaptic D2 receptor. The shapes of the stimulus-response relationship were completely unaffected by age or sex, arguing against presynaptic factors such as reduced release probability. This is also consistent with our results (not shown) using paired trains of stimuli 3 seconds apart to evoke D2-IPSCs in neurons from male mice, which indicated no effect of age (paired-pulse ratio; young: 0.781 ± 0.018, n = 6; old: 0.777 ± 0.011; unpaired t-test p = 0.863). General G protein/GIRK channel activity was also not obviously affected. Bath responses to two concentrations of the GABA_B_ agonist baclofen did not show an effect of age, and the kinetics of the D2-IPSC (which could reflect rapid changes in G protein activation or inactivation) were identical across sex and age. We did observe a reduced response to exogenous dopamine in aged males, indicative of reduced D2 receptor responses, although the effect sizes were small and differences hovered near the edge of statistical significance. Responses to bath applied dopamine seemed to indicate a population of high-responding dopamine neurons in young mice that were not present in old mice, although analysis of peak amplitudes indicated merely a trend toward significance. Interestingly, after several minutes of application currents observed during washout were significantly increased in aged males. This is not likely due to decreased DAT function, as the kinetics of the D2-IPSC are strongly sensitive to DAT-mediated uptake (Beckstead et al., 2004; Branch & Beckstead, 2012) but were unaffected here. This more likely reflects a decrease in desensitization mechanisms following sustained activation. In contrast, analysis of currents elicited by iontophoretic (rapid) application of exogenous dopamine indicated a significantly reduction in aged males. Although the two data sets looked qualitatively similar, the larger effect of age in the local application experiment could be due to the different time course of the respective responses. In bath application experiments, D2 receptor currents peak a few minutes after dopamine application, at a time that receptor activation is already competing with early desensitization. In the iontophoresis experiments, currents peak within a few seconds. It is possible that the reduced desensitization in the aged males following bath application is competing with the lower peak amplitudes, which slightly reduced the effect size when compared to the peak amplitudes obtained following rapid application (either iontophoresis or evoked release of endogenous dopamine). One unfortunate limitation in our interpretation is that there are currently no reliable selective antibodies for measuring total or surface expression of endogenous D2 receptors (Stojanovic et al., 2017). Regardless, the evidence in hand indicates that the most likely explanation for reduced D2-IPSC amplitudes are due to effects at the level of the postsynaptic D2 receptor.

An additional factor that could result in smaller postsynaptic D2 currents is a reduction in somatodendritic morphology in aged males. We previously showed a severely retracted cellular phenotype that worsens with age in the MitoPark mouse model of Parkinson’s disease (Lynch et al., 2018). However, to our knowledge no previous study has analyzed branching parameters in SNc dopamine neurons across the entire lifespan. Our results here show that somatodendritic morphology of dopamine neurons is preserved in aging throughout several dependent measures including multiple levels of dendritic branching. The reduced D2 currents are therefore apparently dependent on effects of the D2 receptor itself, and not merely a product of smaller cells providing less surface area for postsynaptic D2 receptor expression.

In this study, we also explored age effects on CRF-induced enhancement of D2 receptor transmission. D2-IPSCs are subject to multiple forms of pre- (Beckstead et al., 2007) and post-synaptic plasticity including long-term depression induced by the peptide neurotensin (Beckstead & Williams, 2007; Piccart et al., 2015; Tschumi et al., 2022) and a transient enhancement produced by CRF application (Beckstead et al., 2009). The CRF enhancement occurs postsynaptically through activation of CRF1 receptors, and apparently occurs at the level of the GIRK channel, as GABA_B_ signaling is also potentiated. Further, either repeated administration of psychostimulants (cocaine or methamphetamine) or exposure to restraint stress decreasing the magnitude of the response to CRF (Beckstead et al., 2009). Here we show that aging does not affect enhancement of D2-IPSCs produced by bath-application of CRF. Deficits in synaptic plasticity (i.e., long-term potentiation [LTP] of glutamate signaling) have been described with aging that could explain in part some cognitive deficits characteristic of advance age (Bergado & Almaguer, 2002). For example, initial tests for LTP deficits in aged rats (>2-year-old) used acute hippocampal slices and found age-related impairments in the Schaffer/commissural projections from field CA3 to CA1 (Landfield & Lynch, 1977). While it is likely that brain areas exhibit differential sensitivity of synaptic plasticity to advancing age, we have provided evidence that that CRF-induced enhancement of D2-IPSCs and presynaptic short term depression (indicated by paired pulse ratios) in dopamine neurons are not affected by age.

In summary, our results indicate a slight reduction of postsynaptic D2 receptor current amplitudes in SNc dopamine neurons from aged males. Most of the other parameters measured were unaffected, including IPSC kinetics, indicators of presynaptic dopamine release, plasticity induced by CRF, GABA_B_ receptor signaling, and multiple measures of somatodendritic morphology. The overall impression is dendrodendritic transmission in the SNc is predominantly conserved with age.

## Materials and Methods

### Animals

C57BL/6JN male and female mice aged 2-8 months (“young”) and 20-25 months (“old”) were used in this study. Mice were provided by the National Institute on Aging (NIA) aged rodent colony, and these mice or their first and second-generation offspring were used for all experiments. All mice were group-housed in ventilated standard cages with ad libitum access to food and water. Animal rooms were maintained on a reversed 12-hour light/dark cycle (lights off at 9:00 AM) with the temperature held at 26°C. Procedures were in accordance with the Guide for the Care and Use of Laboratory Animals and approved by the Institutional Animal Care and Use Committee at the Oklahoma Medical Research Foundation (OMRF).

### Ex vivo electrophysiology

On the day of the experiment, mice were anesthetized with 2,2,2-Tribromoethanol (0.25 g/kg i.p.) and transcardially perfused with ice-cold carboxygenated (95% O_2_ and 5% CO_2_) cutting solution for 45 seconds. The brains were quickly extracted and placed in a cutting solution containing the following (in mM): 2 KCl, 7 MgCl_2_, 0.5 CaCl_2_, 1.2 NaH_2_PO_4_, 26 NaHCO_3_, 11 D-glucose, and 250 sucrose. Horizontal slices (200 μm) containing the ventral midbrain were obtained using a vibrating microtome (Leica VT1200S). Slices were incubated for 30 min at 33-34°C with carboxygenated artificial cerebrospinal fluid (aCSF) that also contained the NMDA receptor antagonist MK-801 (30 μM). aCSF contained (in mM): 126 NaCl, 2.5 KCl, 1.2 MgCl_2_, 2.4 CaCl_2_, 1.2 NaH_2_PO4, 21.4 NaHCO_3_, and 11.1 D-glucose. Slices were then left to stabilize at room temperature for at least 30 minutes.

Slices were placed in a recording chamber attached to an upright microscope (Nikon Instruments) and maintained at 33-34°C with aCSF perfused at a rate of approximately 2 ml/min. SNc dopaminergic neurons were visually identified based on their location in relation to the midline and the medial terminal nucleus of the accessory optic tract. Dopaminergic neurons from SNc were further identified by their large soma, slow pacemaker firing (< 7 Hz)(Grace & Onn, 1989) with wide extracellular waveforms (>1.1 ms), their characteristic hyperpolarization-activated inward cation current (*I*_H_) of >100 pA (Neuhoff et al., 2002), and ultimately by measuring D2-IPSCs, which have not been reported in other cell types in wild type mice. Recording pipettes (2.0-2.2 M□ resistance) were constructed from thin wall capillaries (World Precision Instruments) with a PC-10 puller (Narishige International). Whole-cell recordings were obtained using an internal solution containing the following (in mM): 115 K-methylsulfate, 20 NaCl, 1.5 MgCl_2_, 10 HEPES-K, 2 ATP, 0.4 GTP, and 10 BAPTA, pH 7.35–7.40, 269 –272 mOsm.

Electrical stimulation was performed with a bipolar platinum electrode with tip separation of 305 μm, placed in the slice ~100 μm caudal to the cell being recorded. The membrane voltage was held at −55 mV and synaptic currents mediated by dopamine could be detected in most of the cells using 5 pulses (0.5 ms duration) applied at 40 Hz once every 60 s. For stimulus-response curve experiments, intensity of the electrical stimulation was applied in an incremental manner (0.02-0.05-0.1-0.15-0.2-0.25-0.3 mA). Iontophoretic pipettes were pulled from thin-wall microelectrodes (resistance ~100 M□), filled with dopamine (1 M), and placed ~20 μm from the cell. A holding current of approximately −3 nA was applied to prevent passive leakage of dopamine, and dopamine was ejected as a cation with a single 1000 ms pulse.

### Drugs

D2 receptor-mediated outward currents were recorded in the presence of the following receptor antagonists (all from Sigma-Aldrich): MK-801 (30 μM added to the recovery bath, NMDA), picrotoxin (100 μM, GABA_A_), CGP 56999A (100 nM, GABA_B_), DNQX (6,7-dinitroquinoxaline-2,3(1H,4H)-dione) (10 μM, AMPA), and hexamethonium (100 μM, nicotinic acetylcholine). For bath application experiments, drug concentrations were as follows: dopamine hydrochloride (Sigma-Aldrich, 100 μM), CRF (Tocris, 100 nM), and baclofen (GABA_B_ agonist, Sigma-Aldrich, 3-30 μM). In iontophoresis experiments, dopamine was dissolved at 1 M. Mg ATP, Na GTP, Na HEPES, K HEPES, and BAPTA were obtained from Sigma-Aldrich. K-methyl sulfate was from MP Biomedicals, LLC.

### Dopamine neuron morphology

To analyze somatodendritic morphology of dopamine neurons, biocytin (Sigma) was added to the internal solution at a final concentration of 0.25% and allowed to diffuse within the cytoplasmic content for 10-12 min with the series resistance stable at < 10 M□. Once cells were filled, and for a maximal preservation of intact soma and dendrites, the glass pipette was slowly retracted horizontally and then vertically until the pipette was outside the slice. A 5-10 min washout with aCSF was allowed to remove any extracellular presence of biocytin molecules that could interfere with the subsequent staining procedure. The 200 μm brain sections were then fixed in 4% paraformaldehyde in phosphate-buffered saline (PBS) for 12 hours and washed three times with PBS at room temperature for at least 30 min. All immunostaining was conducted on freely floating sections, and incubations were conducted with gentle agitation at room temperature. Post-fixed tissue sections were blocked with 5% normal donkey serum (NDS) in PBS with 0.2%Tween 20 (PBST) for 1 hour. Sections were washed at least three times with PBST and incubated with native Streptavidin protein DyLight 488 (1:750 dilution; Abcam, ab134349) in 0.5% fish skin gelatin with PBST for 2 hours. The sections were then subjected to two rinses in PBST followed by two washes in PBS before mounted on gelatin-coated slides, and cover slips were applied with ProLong Gold Antifade Mountant (Thermofisher Scientific, Waltham, MA). Z-stack images were captured using a Zeiss LSM 710 Confocal microscope (Oberkochen, Germany) at 20X magnification with 1 μm stepsize using the Zen software (black edition). Neurites were traced using ImageJ software and the Simple neurite tracer (SNT) plugin. Neurite tracings were used to calculate dendritic measurements and arborization patterns via Sholl and SNT analysis.

### Statistics

Unpaired *Student* t-tests or one-way ANOVA was performed to analyze differences between two or more groups. Two-way ANOVAs were used to analyze differences in current amplitudes over time between groups, along with Holm-Šídák’s multiple comparisons test (Prism 9.3.1, Graphpad). Summary data in graphs is mean ± SEM. Significance was set in all experiments at a p-value < 0.05.

## Acknowledgements

We acknowledge Ben Fowler, Justin Willige, and Julie Crane from the OMRF imaging core for their training and helpful support in the confocal microscopy and in the tracings from dopamine neurons. Funding was provided through National Institute on Aging grants R01 AG052606 and R21 AG072811. Figure 1A was made using Biorender.com.

## Author contributions

MJB was responsible for the study concept and design. ETR, KH, MJB, and SYB contributed to the acquisition of data. All authors contributed to the analysis and interpretation of the findings. MJB and ETR drafted the manuscript with help from KH. MJB, ETR, and KH provided critical revision of the manuscript. All authors reviewed and approved the final version for publication.

## Data availability statement

Data are available from the corresponding author (MJB) upon request.

## Competing Interests Statement

The authors declare no conflicts of interest.

## Notes

### Competing Interest Statement

The authors have declared no competing interest.

## References

Beckstead, M. J., Ford, C. P., Phillips, P. E., & Williams, J. T. (2007). Presynaptic regulation of dendrodendritic dopamine transmission. Eur J Neurosci, 26(6), 1479–1488. https://doi.org/10.1111/j.1460-9568.2007.05775.x

Beckstead, M. J., Gantz, S. C., Ford, C. P., Stenzel-Poore, M. P., Phillips, P. E., Mark, G. P., & Williams, J. T. (2009). CRF enhancement of GIRK channel-mediated transmission in dopamine neurons. Neuropsychopharmacology, 34(8), 1926–1935. https://doi.org/10.1038/npp.2009.25

Beckstead, M. J., Grandy, D. K., Wickman, K., & Williams, J. T. (2004). Vesicular dopamine release elicits an inhibitory postsynaptic current in midbrain dopamine neurons. Neuron, 42(6), 939–946. https://doi.org/10.1016/j.neuron.2004.05.019

Beckstead, M. J., & Williams, J. T. (2007). Long-term depression of a dopamine IPSC. J Neurosci, 27(8), 2074–2080. https://doi.org/10.1523/JNEUROSCI.3251-06.2007

Bergado, J. A., & Almaguer, W. (2002). Aging and synaptic plasticity: a review. Neural Plast, 9(4), 217–232. https://doi.org/10.1155/NP.2002.217

Binley, K. E., Ng, W. S., Tribble, J. R., Song, B., & Morgan, J. E. (2014). Sholl analysis: a quantitative comparison of semi-automated methods. J Neurosci Methods, 225, 65–70. https://doi.org/10.1016/j.jneumeth.2014.01.017

Branch, S. Y., & Beckstead, M. J. (2012). Methamphetamine produces bidirectional, concentration-dependent effects on dopamine neuron excitability and dopamine-mediated synaptic currents. J Neurophysiol, 108(3), 802–809. https://doi.org/10.1152/jn.00094.2012

Branch, S. Y., Sharma, R., & Beckstead, M. J. (2014). Aging decreases L-type calcium channel currents and pacemaker firing fidelity in substantia nigra dopamine neurons. J Neurosci, 34(28), 9310–9318. https://doi.org/10.1523/JNEUROSCI.4228-13.2014

Cohen, F., Alicke, B., Elliott, L. O., Flygare, J. A., Goncharov, T., Keteltas, S. F., Franklin, M. C., Frankovitz, S., Stephan, J. P., Tsui, V., Vucic, D., Wong, H., & Fairbrother, W. J. (2009). Orally bioavailable antagonists of inhibitor of apoptosis proteins based on an azabicyclooctane scaffold. J Med Chem, 52(6), 1723–1730. https://doi.org/10.1021/jm801450c

Colebrooke, R. E., Humby, T., Lynch, P. J., McGowan, D. P., Xia, J., & Emson, P. C. (2006). Age-related decline in striatal dopamine content and motor performance occurs in the absence of nigral cell loss in a genetic mouse model of Parkinson’s disease. Eur J Neurosci, 24(9), 2622–2630. https://doi.org/10.1111/j.1460-9568.2006.05143.x

Courtney, N. A., Mamaligas, A. A., & Ford, C. P. (2012). Species differences in somatodendritic dopamine transmission determine D2-autoreceptor-mediated inhibition of ventral tegmental area neuron firing. J Neurosci, 32(39), 13520–13528. https://doi.org/10.1523/JNEUROSCI.2745-12.2012

Davila, V., Yan, Z., Craciun, L. C., Logothetis, D., & Sulzer, D. (2003). D3 dopamine autoreceptors do not activate G-protein-gated inwardly rectifying potassium channel currents in substantia nigra dopamine neurons. J Neurosci, 23(13), 5693–5697. https://www.ncbi.nlm.nih.gov/pubmed/12843272 https://www.ncbi.nlm.nih.gov/pmc/articles/PMC6741237/pdf/0235693.pdf

Gantz, S. C., Ford, C. P., Neve, K. A., & Williams, J. T. (2011). Loss of Mecp2 in substantia nigra dopamine neurons compromises the nigrostriatal pathway. J Neurosci, 31(35), 12629–12637. https://doi.org/10.1523/JNEUROSCI.0684-11.2011

Grace, A. A., & Onn, S. P. (1989). Morphology and electrophysiological properties of immunocytochemically identified rat dopamine neurons recorded in vitro. J Neurosci, 9(10), 3463–3481. https://www.ncbi.nlm.nih.gov/pubmed/2795134 https://www.jneurosci.org/content/jneuro/9/10/3463.full.pdf

Graves, S. M., Xie, Z., Stout, K. A., Zampese, E., Burbulla, L. F., Shih, J. C., Kondapalli, J., Patriarchi, T., Tian, L., Brichta, L., Greengard, P., Krainc, D., Schumacker, P. T., & Surmeier, D. J. (2020). Dopamine metabolism by a monoamine oxidase mitochondrial shuttle activates the electron transport chain. Nat Neurosci, 23(1), 15–20. https://doi.org/10.1038/s41593-019-0556-3

Groves, P. M., & Linder, J. C. (1983). Dendro-dendritic synapses in substantia nigra: descriptions based on analysis of serial sections. Exp Brain Res, 49(2), 209–217. https://doi.org/10.1007/BF00238581

Guzman, J. N., Sanchez-Padilla, J., Wokosin, D., Kondapalli, J., Ilijic, E., Schumacker, P. T., & Surmeier, D. J. (2010). Oxidant stress evoked by pacemaking in dopaminergic neurons is attenuated by DJ-1. Nature, 468(7324), 696–700. https://doi.org/10.1038/nature09536

Haycock, J. W., Becker, L., Ang, L., Furukawa, Y., Hornykiewicz, O., & Kish, S. J. (2003). Marked disparity between age-related changes in dopamine and other presynaptic dopaminergic markers in human striatum. J Neurochem, 87(3), 574–585. https://doi.org/10.1046/j.1471-4159.2003.02017.x

Hornykiewicz, O. (1975). Parkinson’s disease and its chemotherapy. Biochem Pharmacol, 24(10), 1061–1065. https://doi.org/10.1016/0006-2952(75)90190-2

Howell, R. D., Dominguez-Lopez, S., Ocanas, S. R., Freeman, W. M., & Beckstead, M. J. (2020). Female mice are resilient to age-related decline of substantia nigra dopamine neuron firing parameters. Neurobiol Aging, 95, 195–204. https://doi.org/10.1016/j.neurobiolaging.2020.07.025

Kelly, E. A., & Fudge, J. L. (2018). The neuroanatomic complexity of the CRF and DA systems and their interface: What we still don’t know. Neurosci Biobehav Rev, 90, 247–259. https://doi.org/10.1016/j.neubiorev.2018.04.014

Kim, K. M., Nakajima, Y., & Nakajima, S. (1995). G protein-coupled inward rectifier modulated by dopamine agonists in cultured substantia nigra neurons. Neuroscience, 69(4), 1145–1158. https://doi.org/

Korotkova, T. M., Brown, R. E., Sergeeva, O. A., Ponomarenko, A. A., & Haas, H. L. (2006). Effects of arousal-and feeding-related neuropeptides on dopaminergic and GABAergic neurons in the ventral tegmental area of the rat. Eur J Neurosci, 23(10), 2677–2685. https://doi.org/10.1111/j.1460-9568.2006.04792.x

Kubis, N., Faucheux, B. A., Ransmayr, G., Damier, P., Duyckaerts, C., Henin, D., Forette, B., Le Charpentier, Y., Hauw, J. J., Agid, Y., & Hirsch, E. C. (2000). Preservation of midbrain catecholaminergic neurons in very old human subjects. Brain, 123 (Pt 2), 366–373. https://doi.org/10.1093/brain/123.2.366

Lacey, M. G., Mercuri, N. B., & North, R. A. (1988). On the potassium conductance increase activated by GABAB and dopamine D2 receptors in rat substantia nigra neurones. J Physiol, 401, 437–453. https://doi.org/10.1113/jphysiol.1988.sp017171

Landfield, P. W., & Lynch, G. (1977). Impaired monosynaptic potentiation in in vitro hippocampal slices from aged, memory-deficient rats. J Gerontol, 32(5), 523–533. https://doi.org/10.1093/geronj/32.5.523

Lebowitz, J. J., Trinkle, M., Bunzow, J. R., Balcita-Pedicino, J. J., Hetelekides, S., Robinson, B., De La Torre, S., Aicher, S. A., Sesack, S. R., & Williams, J. T. (2021). Subcellular localization of D2 receptors in the murine substantia nigra. Brain Struct Funct. https://doi.org/10.1007/s00429-021-02432-3

Lodge, D. J., & Grace, A. A. (2005). Acute and chronic corticotropin-releasing factor 1 receptor blockade inhibits cocaine-induced dopamine release: correlation with dopamine neuron activity. J Pharmacol Exp Ther, 314(1), 201–206. https://doi.org/10.1124/jpet.105.084913

Lynch, W. B., Tschumi, C. W., Sharpe, A. L., Branch, S. Y., Chen, C., Ge, G., Li, S., & Beckstead, M. J. (2018). Progressively disrupted somatodendritic morphology in dopamine neurons in a mouse Parkinson’s model. Mov Disord, 33(12), 1928–1937. https://doi.org/10.1002/mds.27541

McGeer, P. L., Itagaki, S., Akiyama, H., & McGeer, E. G. (1988). Rate of cell death in parkinsonism indicates active neuropathological process. Ann Neurol, 24(4), 574–576. https://doi.org/10.1002/ana.410240415

Neuhoff, H., Neu, A., Liss, B., & Roeper, J. (2002). I(h) channels contribute to the different functional properties of identified dopaminergic subpopulations in the midbrain. J Neurosci, 22(4), 1290–1302. https://www.ncbi.nlm.nih.gov/pubmed/11850457 https://www.ncbi.nlm.nih.gov/pmc/articles/PMC6757558/pdf/ns0402001290.pdf

Neve, K. A., Seamans, J. K., & Trantham-Davidson, H. (2004). Dopamine receptor signaling. J Recept Signal Transduct Res, 24(3), 165–205. https://doi.org/10.1081/rrs-200029981

Nimitvilai, S., Arora, D. S., McElvain, M. A., & Brodie, M. S. (2012). Ethanol blocks the reversal of prolonged dopamine inhibition of dopaminergic neurons of the ventral tegmental area. Alcohol Clin Exp Res, 36(11), 1913–1921. https://doi.org/10.1111/j.1530-0277.2012.01814.x

Piccart, E., Courtney, N. A., Branch, S. Y., Ford, C. P., & Beckstead, M. J. (2015). Neurotensin Induces Presynaptic Depression of D2 Dopamine Autoreceptor-Mediated Neurotransmission in Midbrain Dopaminergic Neurons. J Neurosci, 35(31), 11144–11152. https://doi.org/10.1523/JNEUROSCI.3816-14.2015

Piccart, E., Tschumi, C. W., & Beckstead, M. J. (2019). Acute and subchronic PCP attenuate D2 autoreceptor signaling in substantia nigra dopamine neurons. Eur Neuropsychopharmacol, 29(3), 444–449. https://doi.org/10.1016/j.euroneuro.2019.01.108

Pucak, M. L., & Grace, A. A. (1994). Evidence that systemically administered dopamine antagonists activate dopamine neuron firing primarily by blockade of somatodendritic autoreceptors. J Pharmacol Exp Ther, 271(3), 1181–1192. https://www.ncbi.nlm.nih.gov/pubmed/7996424

Reeve, A., Simcox, E., & Turnbull, D. (2014). Ageing and Parkinson’s disease: why is advancing age the biggest risk factor? Ageing Res Rev, 14, 19–30. https://doi.org/10.1016/j.arr.2014.01.004

Robinson, B. G., Cai, X., Wang, J., Bunzow, J. R., Williams, J. T., & Kaeser, P. S. (2019). RIM is essential for stimulated but not spontaneous somatodendritic dopamine release in the midbrain. Elife, 8. https://doi.org/10.7554/eLife.47972

Sauvage, M., & Steckler, T. (2001). Detection of corticotropin-releasing hormone receptor 1 immunoreactivity in cholinergic, dopaminergic and noradrenergic neurons of the murine basal forebrain and brainstem nuclei--potential implication for arousal and attention. Neuroscience, 104(3), 643–652. https://doi.org/10.1016/s0306-4522(01)00137-3

Schultz, W. (2007). Multiple dopamine functions at different time courses. Annu Rev Neurosci, 30, 259–288. https://doi.org/10.1146/annurev.neuro.28.061604.135722

Seidler, R. D., Bernard, J. A., Burutolu, T. B., Fling, B. W., Gordon, M. T., Gwin, J. T., Kwak, Y., & Lipps, D. B. (2010). Motor control and aging: links to age-related brain structural, functional, and biochemical effects. Neurosci Biobehav Rev, 34(5), 721–733. https://doi.org/10.1016/j.neubiorev.2009.10.005

Sharpe, A. L., Varela, E., Bettinger, L., & Beckstead, M. J. (2014). Methamphetamine self-administration in mice decreases GIRK channel-mediated currents in midbrain dopamine neurons. Int J Neuropsychopharmacol, 18(5). https://doi.org/10.1093/ijnp/pyu073

Shin, J., Kovacheva, L., Thomas, D., Stojanovic, S., Knowlton, C. J., Mankel, J., Boehm, J., Farassat, N., Paladini, C., Striessnig, J., Canavier, C. C., Geisslinger, G., & Roeper, J. (2022). Cav1.3 calcium channels are full-range linear amplifiers of firing frequencies in lateral DA SN neurons. Sci Adv, 8(23), eabm4560. https://doi.org/10.1126/sciadv.abm4560

Stojanovic, T., Orlova, M., Sialana, F. J., Hoger, H., Stuchlik, S., Milenkovic, I., Aradska, J., & Lubec, G. (2017). Validation of dopamine receptor DRD1 and DRD2 antibodies using receptor deficient mice. Amino Acids, 49(6), 1101–1109. https://doi.org/10.1007/s00726-017-2408-3

Surmeier, D. J., Guzman, J. N., Sanchez-Padilla, J., & Goldberg, J. A. (2011). The origins of oxidant stress in Parkinson’s disease and therapeutic strategies. Antioxid Redox Signal, 14(7), 1289–1301. https://doi.org/10.1089/ars.2010.3521

Tagliaferro, P., & Morales, M. (2008). Synapses between corticotropin-releasing factor-containing axon terminals and dopaminergic neurons in the ventral tegmental area are predominantly glutamatergic. J Comp Neurol, 506(4), 616–626. https://doi.org/10.1002/cne.21576

Taylor, W. D., Zald, D. H., Felger, J. C., Christman, S., Claassen, D. O., Horga, G., Miller, J. M., Gifford, K., Rogers, B., Szymkowicz, S. M., & Rutherford, B. R. (2022). Influences of dopaminergic system dysfunction on late-life depression. Mol Psychiatry, 27(1), 180–191. https://doi.org/10.1038/s41380-021-01265-0

Tschumi, C. W., Blankenship, H. E., Sharma, R., Lynch, W. B., & Beckstead, M. J. (2022). Neurotensin Release from Dopamine Neurons Drives Long-Term Depression of Substantia Nigra Dopamine Signaling. J Neurosci. https://doi.org/10.1523/JNEUROSCI.1395-20.2022

Ungless, M. A., Singh, V., Crowder, T. L., Yaka, R., Ron, D., & Bonci, A. (2003). Corticotropin-releasing factor requires CRF binding protein to potentiate NMDA receptors via CRF receptor 2 in dopamine neurons. Neuron, 39(3), 401–407. https://doi.org/10.1016/s0896-6273(03)00461-6

Wanat, M. J., Hopf, F. W., Stuber, G. D., Phillips, P. E., & Bonci, A. (2008). Corticotropin-releasing factor increases mouse ventral tegmental area dopamine neuron firing through a protein kinase C-dependent enhancement of Ih. J Physiol, 586(8), 2157–2170. https://doi.org/10.1113/jphysiol.2007.150078

Wang, T. F., Wu, S. Y., Pan, B. S., Tsai, S. F., & Kuo, Y. M. (2022). Inhibition of Nigral Microglial Activation Reduces Age-Related Loss of Dopaminergic Neurons and Motor Deficits. Cells, 11(3). https://doi.org/10.3390/cells11030481

Wooten, G. F. C., L. J.; Bovbjerg, V.E.; Lee, J.K.; Patrie, J. (2004). Are men at greater risk for Parkinson’s disease than women? J Neurol Neurosurg Psychiatry, 75, 637–639. https://doi.org/10.1136/jnnp.2003.020982

Yin, H. H., Ostlund, S. B., & Balleine, B. W. (2008). Reward-guided learning beyond dopamine in the nucleus accumbens: the integrative functions of cortico-basal ganglia networks. Eur J Neurosci, 28(8), 1437–1448. https://doi.org/10.1111/j.1460-9568.2008.06422.x

